# Hydropathy patterning complements charge patterning to describe conformational preferences of disordered proteins

**DOI:** 10.1101/2020.01.25.919498

**Authors:** Wenwei Zheng, Gregory Dignon, Matthew Brown, Young C. Kim, Jeetain Mittal

## Abstract

Understanding the conformational ensemble of an intrinsically disordered protein (IDP) is of great interest due to its relevance to critical intracellular functions and diseases. It is now well established that the polymer scaling behavior can provide a great deal of information about the conformational properties as well as liquid-liquid phase separation of an IDP. It is, therefore, extremely desirable to be able to predict an IDP’s scaling behavior from the protein sequence itself. The work in this direction so far has focused on highly charged proteins and how charge patterning can perturb their structural properties. As naturally occurring IDPs are composed of a significant fraction of uncharged amino acids, the rules based on charge content and patterning are only partially helpful in solving the problem. Here, we propose a new order parameter, sequence hydropathy decoration (SHD), which can provide a near quantitative understanding of scaling and structural properties of IDPs devoid of charged residues. We combine this with a charge patterning parameter, sequence charge decoration (SCD), to obtain a general equation, parameterized from extensive coarse-grained simulation data, for predicting protein dimensions from the sequence. We finally test this equation against available experimental data and find a semi-quantitative match in predicting the scaling behavior. We also provide guidance on how to extend this approach to experimental data, which should be feasible in the near future.

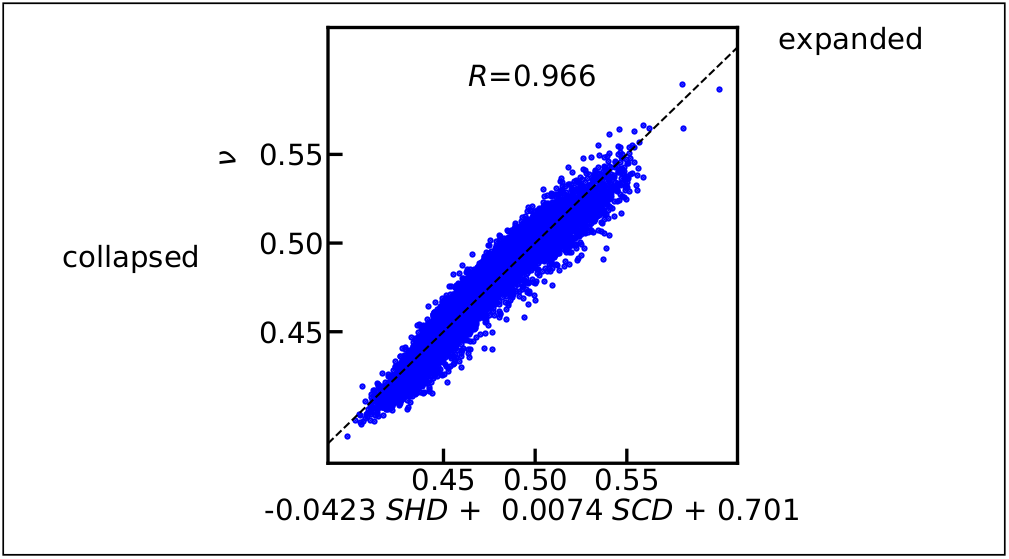
Graphical TOC Entry.

---

Intrinsically disordered proteins (IDPs) are of great interest in biology due to their involvement in important intracellular functions and pathological diseases.^1–4^ These proteins lack a well-defined three-dimensional structure and are more appropriately described by a conformational ensemble in contrast to folded proteins with a single folded structure. ^5,6^ It is, therefore, nontrivial to study IDPs via the traditional structure-function relationship considering the heterogeneous nature of an IDP conformational ensemble. However, one still expects that the function of an IDP is determined by its sequence,^7,8^ as observed in numerous cases.^9–13^ It is important to identify sequence-dependent structural ensemble features capable of bridging the gap between sequence and function of an IDP, so that the structure-function paradigm can still be applied to IDPs.^14^

A variety of fundamental features ranging from average residue-level structural details to overall protein dimensions can be important for characterizing the conformational properties of an IDP. Nuclear magnetic resonance (NMR) experiments alone or coupled with all-atom simulations arguably provide the most detailed information on residual secondary structure properties and inter-residue interactions.^5,12,15–19^ These data can help generate knowledge of how specific amino acids^20^ and interactions between pairs of amino acid types may dictate the IDP properties.^19,21,22^ Such empirical rules are significant for understanding the behavior of low complexity IDP sequences that are composed of only a few types of amino acids.^23^ On the other hand, small-angle X-ray scattering (SAXS)^24^ and Förster resonance energy transfer (FRET)^25^ experiments provide estimates of global protein dimensions such as the radius of gyration (*R_g_*). The interpretation of these experiments in terms of the polymer scaling behavior of proteins is helpful in applying existing analytical theories.^26–32^

Polymer scaling exponent (*ν*) is commonly used to characterize the relationship of the polymer size in solution with its chain length *N* as *R_g_ ∝ N^ν^*. This variable also provides information on the solvent quality in terms of good, bad, or ideal solvent.^26,27^ Despite the sequence heterogeneity of IDPs that contain twenty naturally occurring amino acids (and possibly many other non-canonical amino acids), there is increasing evidence that a single *ν* value may be used to characterize the conformational properties of disordered proteins. For instance, we recently showed that a *ν*-dependent distance distribution function based on a self-avoiding random walk model could help interpret experimental data from FRET^33^ and SAXS.^30^ All-atom simulation data for more than 30 protein sequences further strengthened the understanding that IDPs behave as an ideal chain (*ν*=0.5) in aqueous solution and as an excluded volume chain (*ν ≈* 0.58) in high denaturant concentrations.^20,28,34–36^ We also found that the *θ*-temperature (*T_θ_*) of a single chain, the temperature at which *ν*=0.5, is strongly correlated with the critical temperature (*T_c_*) of liquid-liquid phase separation (LLPS) of disordered proteins.^37,38^ This relationship provides a rapid method for approximating the behavior of IDPs in the context of LLPS, and aided in the development of a novel temperature-dependent interaction potential that explained upper- and lower critical solution temperature phase transitions based on temperature-dependent solvent-mediated interactions.^39^

Given the role of polymer scaling properties in dictating the conformational behavior of IDPs, there have been significant efforts to predict *ν* as a function of the protein sequence or order parameters representing important sequence characteristics. A protein’s net charge and average hydropathy can help distinguish foldable sequences from disordered ones,^40^ but it is essential also to consider other features such as the fraction of charged residues, and their patterning within the chain.^41–44^ It is likely that the patterning of all amino acids, including uncharged ones, can contribute to the behavior of IDPs. Up to this point, however, this has not been studied in the context of 20 different amino acid types, even though it could be expected to be quite crucial, particularly for natural IDP sequences that contain a significant fraction of uncharged residues.^45–47^

In this work, we use our recently developed coarse-grained model of IDPs^48^ to study the role of sequence patterning of uncharged residues in an extensive data set containing 5130 sequences. As expected, average hydropathy alone is not able to explain the sequence-dependent scaling behavior well, which leads us to develop a new sequence hydropathy decoration (SHD) parameter motivated by the extensively used sequence charge decoration (SCD) parameter.^43^ The new SHD parameter performs much better than the commonly used average hydropathy in reproducing the polymer scaling properties of these sequences, demonstrating the importance of patterning of twenty amino acids, beyond just charge patterning, in characterizing the size of the IDPs. We further find that the combination of SHD and SCD can capture the scaling properties of a more extensive data set (10260 sequences) containing all twenty amino acids remarkably well. Based on these results, we propose that a combination of SHD and SCD can be used to rapidly predict the scaling behavior of the disordered proteins and pave the way for high throughput screening of disordered sequences before wet lab investigation. We demonstrate this in the context of disordered protein sequences using the Disprot (disordered) database^23^ and Top8000 (folded)^49^ database as a control. We finally test predictions from our equation against existing experimental measurements of the size of several disordered proteins.

## Computational estimation of polymer scaling exponent *ν* of IDPs

The advantage of using *ν* as opposed to *R_g_* to characterize a protein’s size is to eliminate the chain length dependence and to provide meaningful information on the solvent quality that can be useful in predicting protein LLPS.^37^ We have recently shown that *R_g_* of a single protein can be used to estimate the scaling exponent (*ν_Rg_*):^33,50,51^

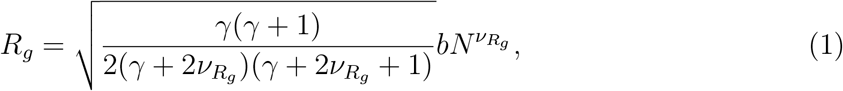

where *γ ≈*= 1.1615,^52^ *b* = 0.55 nm,^28,33^ and *N* is the number of peptide bonds (i.e., one less than the number of residues). Alternatively, when analyzing molecular simulation data, *ν_fit_* can be obtained using the following equation, which is based on the mean intrachain distance *R_i,j_* as a function of sequence separation *|i − j|*,^41,53^

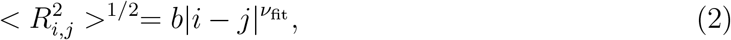

where *b* = 0.55 nm as in the Eq. 1. In practice, Eq. 1 provides a more convenient way of estimating *ν* from simulation data set, and more importantly, from experimentally determined *R_g_*. However, its validity over the whole range of compactness of IDPs has not yet been established. Thus, it is important for us to test whether these two definitions give consistent results in predicting *ν*.

For this purpose, we generated a large set of 10,260 random protein sequences, having chain lengths in the range from 30 to 200 residues, and with amino acid probabilities set equal to their relative abundance in natural IDPs (see database A in Fig. S1).^54^ We then conducted simulations of all of these sequences using our recently developed coarse-grained (CG) model, which represents each amino acid as a single interaction site (See Supporting Methods and ref.^48^). We find that the two methods of calculating *ν* are highly correlated, as shown in Fig. 1. Slight deviations are observed at low and high *ν* values, which suggests that the two methods will yield somewhat different scaling exponents. We asked if these deviations are related to an easily identifiable source in terms of protein’s sequence properties, such as the chain length. As shown in Fig. S2, chain length does seem to cause some discernible differences in the *ν* estimates based on the two methods. Further analysis suggests that for low *ν* values, the *ν*_fit_ estimate may not be appropriate as the intrachain distance fits are not optimal over the whole range of sequence separation (see Fig. S3). For higher *ν* values, one may have to use a different prefactor *b* while using the intrachain distance fits to obtain *ν_fit_*. The parameters used in Eq. 1 (i.e. *γ* and *b*) are almost optimal for minimizing the averaging deviations between *ν*_fit_ and *ν*_Rg_ for the whole range of *ν* values as shown in Fig. S4. For simplicity and keeping in mind that the relative errors across the whole database are mostly less than 5%, we suggest using Eq. 1 to reliably estimate a protein’s scaling behavior in this and future studies.

**Figure 1:**
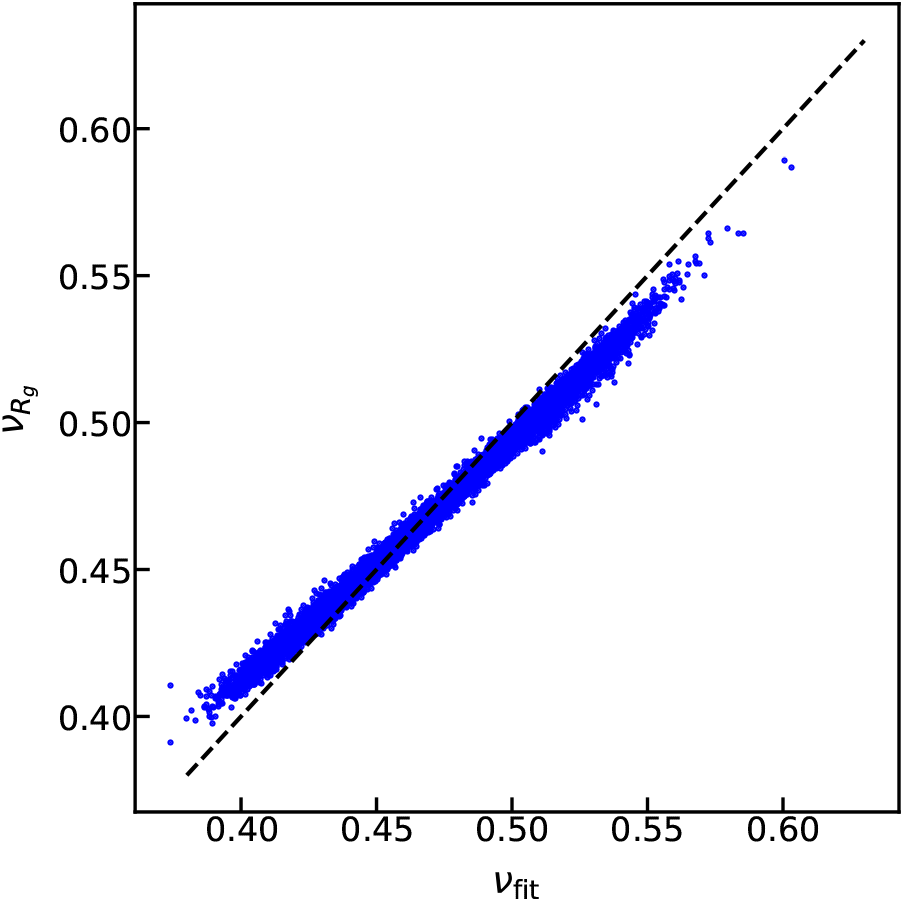
Comparison between the polymer scaling exponents obtained by fitting intramolecular distances (*ν*_fit_) or by using Eq. 1(*ν*_Rg_).

## Sequence hydropathy decoration (SHD) parameter describes properties of uncharged IDPs

Significant previous work has already highlighted the role of sequence charge patterning on the properties of IDPs and important order parameters, such as sequence charge decoration (SCD) and *κ*, are available to describe such effects.^42–44,55,56^ Given the success of such strategies, we focus on developing a descriptor for the patterning of uncharged residues. Here we start with an amino-acid specific hydropathy value (*λ*),^57^ normalized to a value between 0 and 1, as the relevant feature to describe or predict a protein’s compactness, and equivalently the polymer scaling properties. To isolate the effects of non-electrostatic interactions on chain dimensions, we generated an additional protein database of 5130 sequences in which the charged residues are not incorporated, and otherwise preserving the relative abundance of the uncharged amino acids (database B in Fig. S1). Interestingly, in the absence of charged residues, the average hydropathy (*< λ >*) of the sequence does not predict its *ν* very well (Fig. 2A) with only a weak trend showing that higher *< λ >* values are generally more compact. Thus, additional considerations are needed in order to better capture the sequence-dependent properties originating from the patterning of uncharged residues.

**Figure 2:**
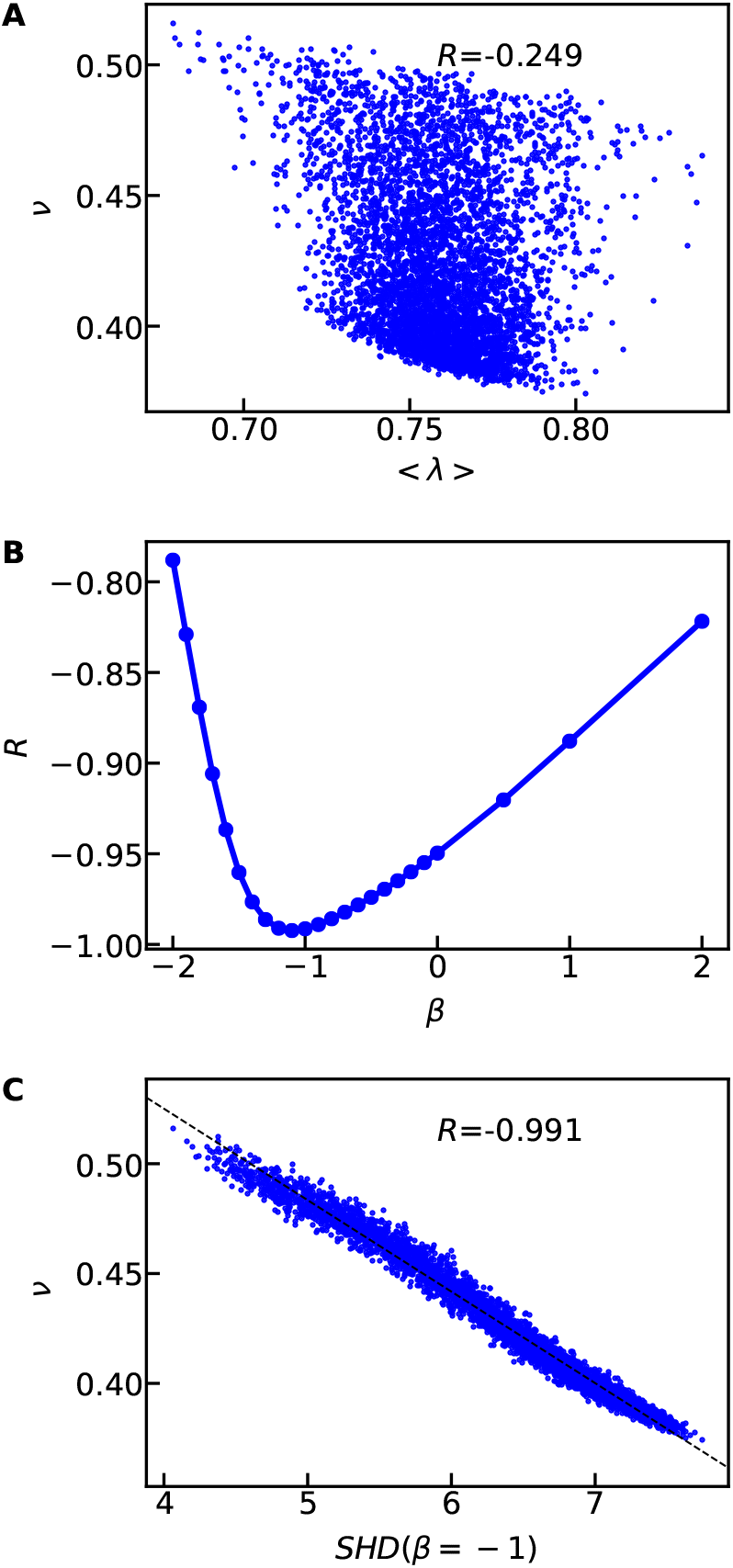
A) Using the mean hydropathy (*< λ >*) to capture the scaling exponents of random uncharged sequences. B) Pearson correlation coefficient between *SHD* and *ν* when varying *β* in Eq. 4. C) Using the hydropathy patterning parameter *SHD* with *β* = *−*1 to capture the scaling exponents. The dashed lines show the linear fitting between *SHD* and *ν* and the legends show the Pearson correlation coefficients.

Motivated by the success of the *SCD* parameter in capturing the charge patterning effects on protein behavior,^37,43,58,59^ we propose a similar strategy to describe effects of hydropathy patterning, terming the new parameter sequence hydropathy decoration (*SHD*). To derive *SHD*, we adopt the theoretical approach presented in detail by Sawle and Ghosh^43^ for charged polymers. The excluded volume contribution to the end-to-end distance of a polymer can be expressed as Ω (see Eq. 13 of Ref.^43^)

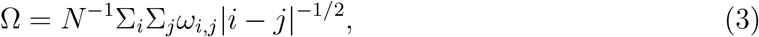

where *ω_i,j_* is the excluded volume interaction parameter between the two residues *i* and *j*. In our IDP model, such interactions arise from the short-range pairwise interactions with the interaction strength of (*λ_i_* + *λ_j_*) for a pair of residues *i* and *j*. The simplest assumption, namely, *ω_i,j_* = *λ_i_* + *λ_j_*, then leads to the following equation,

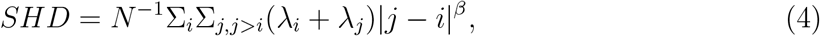

in which *β* = *−*1/2 accounts for the contribution of sequence separation. This choice of the exponent shows much greater predictive ability than just *< λ >* with a much higher negative Pearson correlation coefficient (−0.974) between *ν* and *SHD* values (see Fig. S5). In general, a high value of *SHD* corresponds to sequences with higher *< λ >*, and more clustering (patchiness) of hydrophobic residues together, resulting in a more collapsed conformation.

Next, we asked if the correlation can be improved by changing the *β* value to account for factors not considered in Eq. 4. We find that *SHD* provides the best description of our data for the *β* value *≈ −*1 as shown in Fig. 2B and Fig. S5. This *β* value suggests that *ω_i,j_* can also be sequence separation dependent. Since the two residues will be less likely in contact when the average distance between them becomes large, we can assume *ω_i,j_* is inversely proportional to the average distance between the two residues. We can, therefore obtain

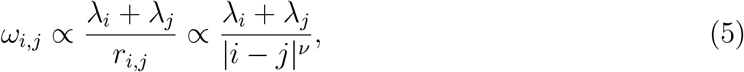

where the last part of the equation makes use of the expected distance dependence from the polymer scaling law. This leads to *β* = *−ν −* 1/2, where for IDPs with 0.45 *≤ ν ≤* 0.6, one gets *−*1.1 *≤ β ≤ −*0.95, which is in excellent agreement with *β* obtained from the simulation data (see Fig. 2C and Fig. S4). We also find that this empirical approach to obtaining the sequence separation exponent (*β* in Eq. 4) also recovers the known exponent value for *SCD* (0.5 as derived by Sawle and Ghosh;^43^ Fig. S6). For simplicity, we set *β* = *−*1 for calculating *SHD* resulting in good correlation with *ν* from simulations (*R* = *−*0.991) as shown in Fig. 2C. This is a huge improvement over the commonly used average hydropathy (*R* = *−*0.249). The observed dependence of hydropathy patterning on the sequence separation is weaker (*β* = *−*1) as compared to the charge patterning (*β* = 0.5), which could be expected considering their differences in interaction range. Thus we find that by developing the *SHD* parameter, we are able to make accurate predictions of IDP scaling behavior simply from the sequence, assuming the absence of charged amino acids.

## Predicting scaling behavior from sequence descriptors

We then investigate how *SHD* compares with the other sequence descriptors (Table S1) to characterize *ν*, particularly in the case of sequences containing charged residues as well. We first look at the correlation between all sequence descriptors (independent of each other) and *ν* (Fig. S7A) and find that the most representative descriptors are *SHD*, *< λ >*, and *SCD*.The importance of *< λ >* is consistent with previous work showing that it can be used to categorize disordered proteins.^40^ However *SHD* and *SCD* stand out, which is probably due to the detailed nature of these two descriptors, accounting not only for the average value, but also patterning.

We expect that at least two sequence descriptors – one relevant to the amino acid charges and the other describing amino acid hydropathy – will be needed to describe the properties of IDPs. To test whether these metrics can work cooperatively to predict *ν*, we scan every pair of sequence descriptors using multilinear regression. In Fig. 3A, we show the Pearson correlation coefficients between the predicted *ν* from each pair of sequence descriptors and the simulated *ν* using the sequence database A (which contains charged amino acids; Fig. S1). While *< λ >* scored higher than *SCD* in the single parameter regression (Fig. S7A), what it provides is redundant when used with *SHD*, so the combination of *< λ >* and SHD does not significantly improve predictions over just *SHD*. The pairing of *SHD* and *SCD* results in the highest Pearson correlation coefficient (0.966) between the predicted and simulated *ν*. The multilinear equation for using *SHD* and *SCD* to predict *ν* and the corresponding *R_g_* (using Eq. 1) is,

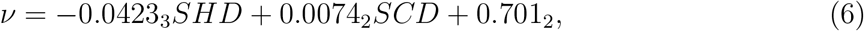

in which the subscripts show the errors of the last digit. The errors were estimated by randomly splitting the sequences into five groups for obtaining the standard deviation of the fitting parameters and repeating the random selection 100 times for averaging the errors. A linear regression using only *SHD* gives similar fitting parameters in comparison to Eq. 6 for the *SHD* prefactor and the constant term (Fig. S7B).

**Figure 3:**
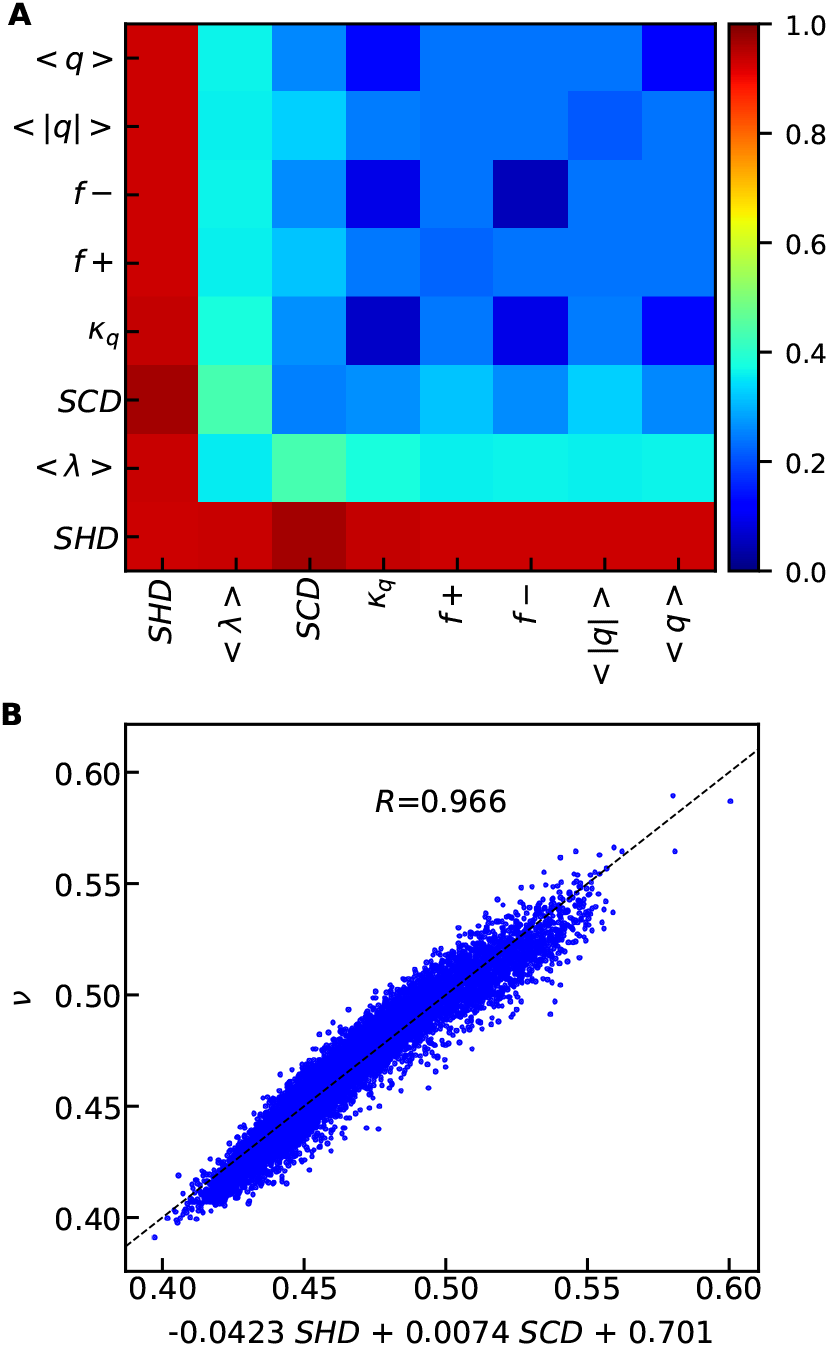
Capturing the scaling exponents (*ν*) using linear models of two sequence descriptors. A) Pearson correlation coefficients between the linearly modelled and simulated *ν*. B) The model based on the best pair of sequence descriptors with the linear equation shown in labels of x-axis.

Eq. 6 should also work for IDP sequences without any charged amino acids since *SCD* goes to zero. However, when the fraction of charged amino acids (*< |q| >*) increases, we expect that the contribution of *SCD* to the compactness of the chain should also become more important. This can be verified by performing the multilinear regression for subsets of our IDP sequence database with different values of *< |q| >*. As shown in Fig. S8, we find that the three fitting parameters do not change that much for *< |q| >* values from 0.2 to 0.3. The *SCD* prefactor starts to increase when *< |q| >* is greater than 0.3, as expected. We further assess relative effectiveness of different sequence descriptors and *ν* for different ranges of *< |q| >* values (Fig. S9). We see that the charge patterning descriptors, *SCD* and *κ*, become increasingly important in determining the chain properties with increasing charge content (*< |q| >*). This is also consistent with previous literature that for sequences with all charged amino acids, charge patterning parameters are most important in characterizing the dimension of the chain.^42,43,60^ However, because a large fraction of disordered proteins have *< |q| >* smaller than 0.3, one needs for the role of hydropathy patterning, which we propose can be accomplished using *SHD*.

## Experimental verification of the simplified equation based on *SHD* and *SCD*

Since we find that *SHD* and *SCD* together can be used to predict *ν* from the simulation data set based on a simplified CG model of IDPs, we would like to test the model’s transferability by using known disordered and folded protein sequences. We select the disordered protein sequences from the Disprot database,^23^ excluding sequences having the disordered region shorter than 30 residues, as the polymer scaling law description may not work well for shorter chain lengths. For each sequence, only the longest disordered region is selected resulting in a total of 557 disordered protein sequences. We select folded protein sequences from a protein database Top8000,^49^ in which the structure of every sequence has been solved with a high-resolution experimental method. We exclude the sequences in the database for which multiple chains are present, resulting in a total of 2360 folded protein sequences. We show in Fig. 4 that Eq. 6, using a combination of *SHD* and *SCD* obtains an average *ν* value close to 0.5 for the disordered proteins, consistent with previous knowledge in the field that disordered proteins in aqueous conditions behave similar to a Gaussian chain. As a control, the *ν* values for the folded proteins predicted using the model are generally smaller.

**Figure 4:**
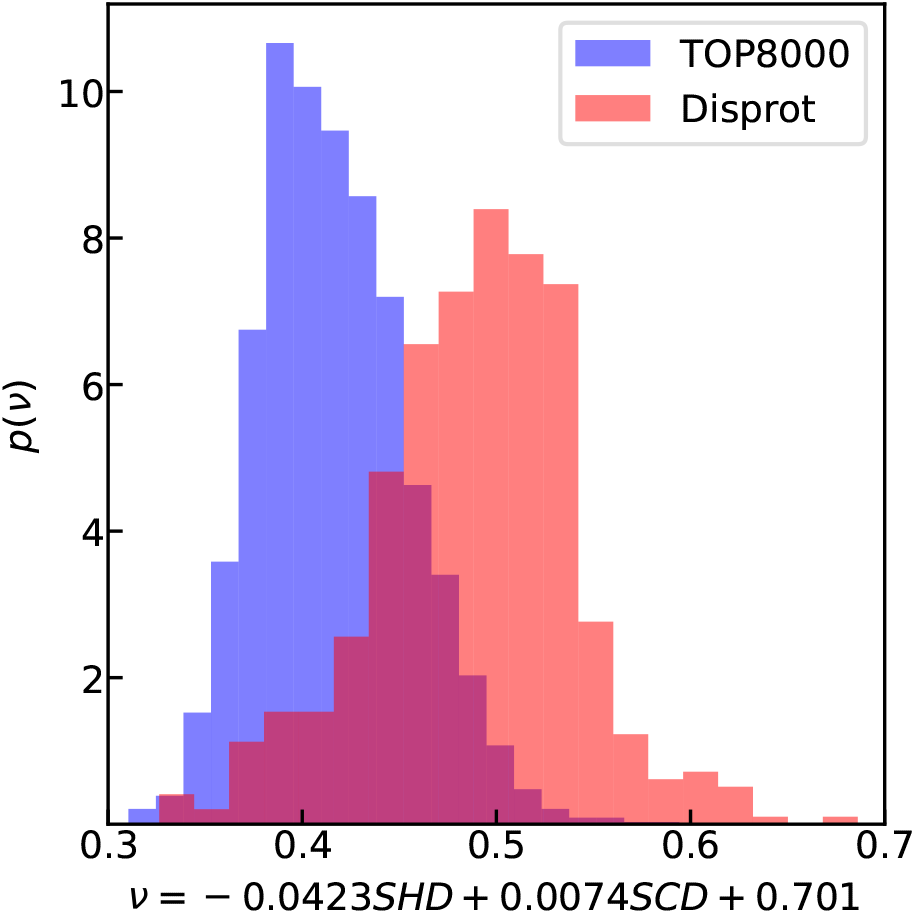
Using *SHD* and *SCD* to predict *ν* of disordered sequences from Disprot database (red).^23^ The folded sequences using TOP8000 database^49^ are shown in blue as a control.

It is likely that the use of a simple CG model to parameterize Eq. 6 would introduce errors based on the limitations of the model. Thus, we would like to further ask if one can directly use experimental data to parameterize Eq. 6 and how many sequences with experimentally determined *R_g_* are necessary to obtain such an optimal predictive equation. We estimate the number of experimental sequences needed by splitting our computational database into two sets–a training set for fitting *ν* and a test set for checking the accuracy of the resulting model. For consistency, the number of sequences in the test set is fixed at a quarter of the total number of sequences (2565 of 10260 sequences) in database A (Fig. S1) while reducing the number of sequences in the training set. The process of randomly selecting the sequences to form the training and test sets is repeated 100 times to obtain the averaging errors of the model. We observe a typical L-shape plot for the relative errors as a function of the number of sequences (Fig. S10), which suggests that about 100 sequences will be sufficient to obtain an accurate predictive model. We expect that the actual number of sequences may differ as the estimate above is based on randomly generated protein sequences that may not capture the diversity of naturally occurring protein sequences, which are not completely random due to pressure from natural selection. Still, one expects the number of protein sequences necessary to obtain an experimentally validated predictive model to be within reach, especially if these sequences are carefully designed. Interestingly, the relative error is still quite reasonable, even for considerably smaller training sets (Fig. S10). Data-driven methods may prove to be quite helpful in this regard.

We then test currently available data on IDPs from the existing literature to validate our predictions. There has been an increasing number of experimental measurements on the compactness of disordered proteins, using either FRET or SAXS. However, there is clear difficulty of directly using the available data for parameterizing an empirical equation due to difficulties in interpreting experimental measurements. Recent work has shown that interpreting *R_g_* or *ν* from FRET or SAXS experiments is not trivial due to the heterogeneous conformations disordered proteins can adopt.^61,62^ FRET experiments tended to underestimate the *R_g_* due to the assumptions about underlying distance distribution, whereas SAXS experiments overestimated the *R_g_* due to a non-linearity in the Guinier plot.^30,61^ We have identified a list of disordered sequences from three publications,,^28,61,63^ two of which contain consistent FRET and SAXS measurements.^61,63^ We reanalyzed these data using our recently published method: SAW-*ν*,^33^ in which a *ν*-dependent distance distribution function is used to adapt the variation of chain dimension (see details in Table S2). As shown in Fig. 5, predicted *ν* for these proteins using Eq. 6 are in reasonable agreement with the experimental *ν* values that are derived using the SAW-*ν* model. It is even more remarkable if we consider the simplicity of our linear model combined with the lack of parameterization to account for different ionic strengths in these experiments. In the future, experiments conducted at similar conditions may be used as inputs to our framework, so that a model solely based on experimental data can be built for directly predicting the IDP conformations at those conditions.

**Figure 5:**
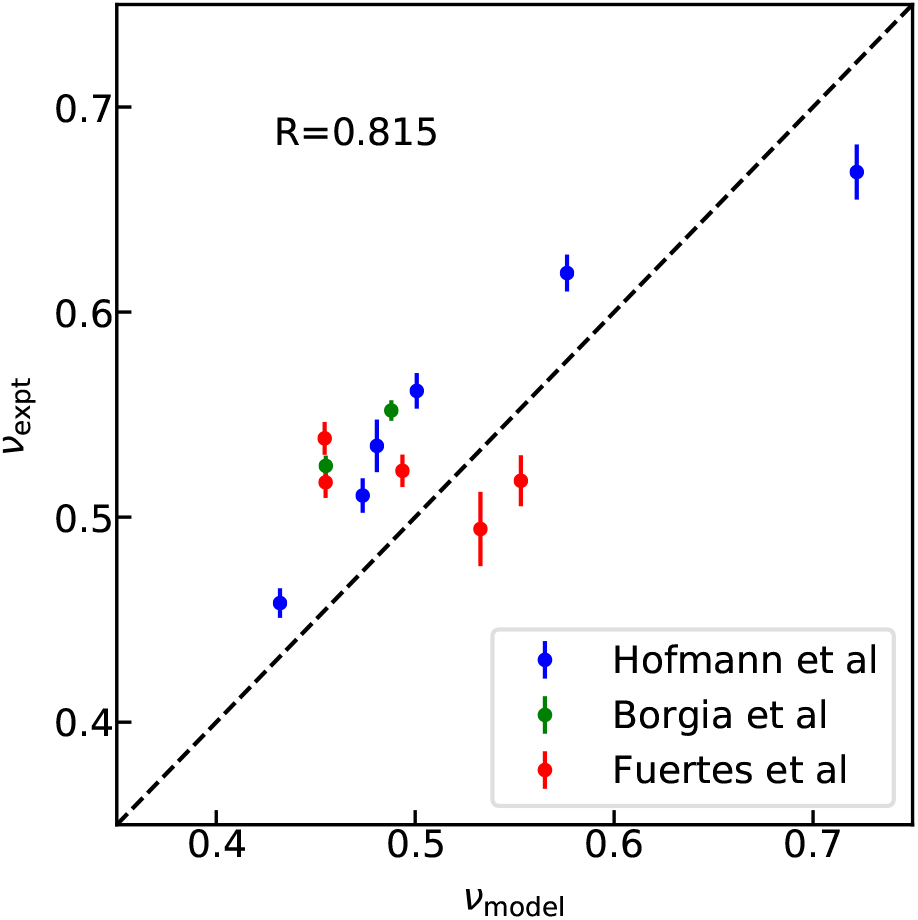
Comparison between the *R_g_* (A) and *ν* (B) from linear model using *SHD* and *SCD* and from FRET and SAXS experiments.

## Conclusion

Intrinsically disordered proteins perform a myriad of biological functions and are also involved in several debilitating disease conditions, but the sequence-structure-(mis)function relationships of these proteins are not well understood. The first step in developing such relationships is to understand better how the conformational preferences of disordered proteins originate from their sequence. Previous work has highlighted the role of charge content and patterning in developing sequence-structure relationships of highly charged proteins to capture the effects of electrostatic interactions. There has been relatively little progress in accounting for the role of other types of interactions such as van der Waals, hydrogen bonding, through which uncharged amino acids interact. We propose that the amino acid hydropathy value can serve as a useful proxy to capture the average interactions of different amino acids, and how it affects the protein dimensions as part of a chain. To describe the presence and arrangement of amino acids with varying values of hydropathy, we propose a sequence hydropathy decoration parameter that can quantitatively capture the sequence-structure relationship for an extensive set of disordered proteins (lacking charged residues) simulated using a coarse-grained model. We combine this new parameter with the existing sequence charge decoration parameter to quantitatively predict protein dimensions simply based on the protein sequence. We anticipate that the predictive equation can serve as a quick screening tool to design new protein sequences with tunable properties as well as allow for future rapid optimization of coarse-grained models to better reproduce experimental results. Most importantly, we can already describe the scaling behavior of many proteins for which experimental data are available from single-molecule FRET and SAXS. This work should significantly contribute towards a quantitative understanding of a disordered proteins’ sequence-structure relationship, which we expect to apply to a better understanding of protein function as well.

## Acknowledgement

This work was partially supported by the National Institutes of Health grants R01GM118530 and R01GM120537. Use of the high-performance computing capabilities of the Extreme Science and Engineering Discovery Environment (XSEDE), which is supported by the National Science Foundation, project no. TG-MCB120014, is also gratefully acknowledged. W.Z. acknowledges the Research Computing at Arizona State University for providing HPC and storage that have contributed to the research results reported within this paper. Y.C.K is supported by the Office of Naval Research via the U.S. Naval Research Laboratory base program.

## Supporting Information Available

Supporting methods, figures and tables.

## Supporting Information

### 1. Supplementary Methods

#### Molecular simulation methods

The simulations of sequences in the database (Fig. S1) were conducted using the HOOMD-Blue v2.1.5.[1] For each sequence, the simulation were run for 500 ns using a Langevin thermostat with a friction coefficient of 1 ps*^−^*^1^, and the first 100 ns was dumped as equilibration. A time step of 10 fs and a temperature of 300 K were used for all the simulations. The ionic strength is set to be 100 mM and characterized using Debye-Hückle electrostatic screening [2]. There are three types of interactions, including bonded, electrostatic and short-range pairwise interaction term characterized by amino acid hydrophobicity.[3] We refer the readers to the literature of our HPS model for the details of the coarse-grained model.[4]

### 2. Supplementary Figures

**Figure S1.**
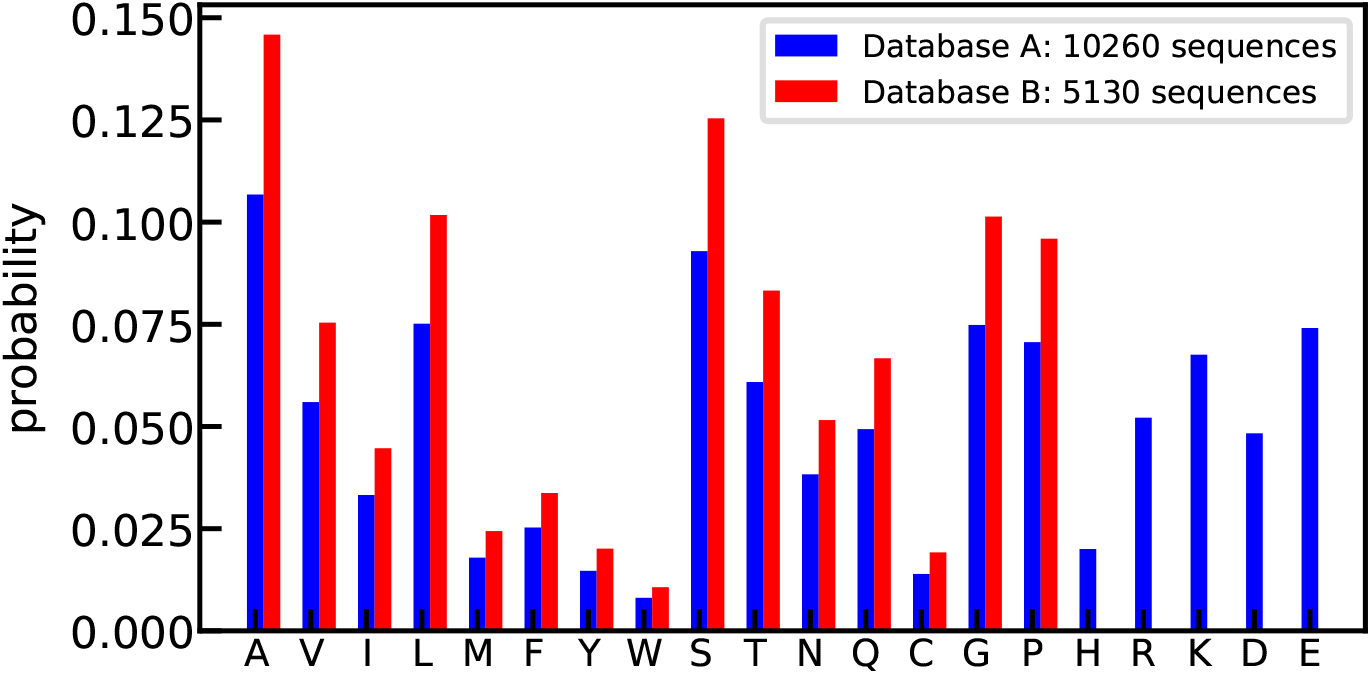
Amino acid probabilities for the randomly generated sequences. Database A contains 10260 charged random sequences (blue), in which there are 60 sequences for each of the chain length from 30 to 200, and database B contains 5130 random sequences without charged amino acids (red), in which there are 30 sequences for each of the chain length.

**Figure S2.**
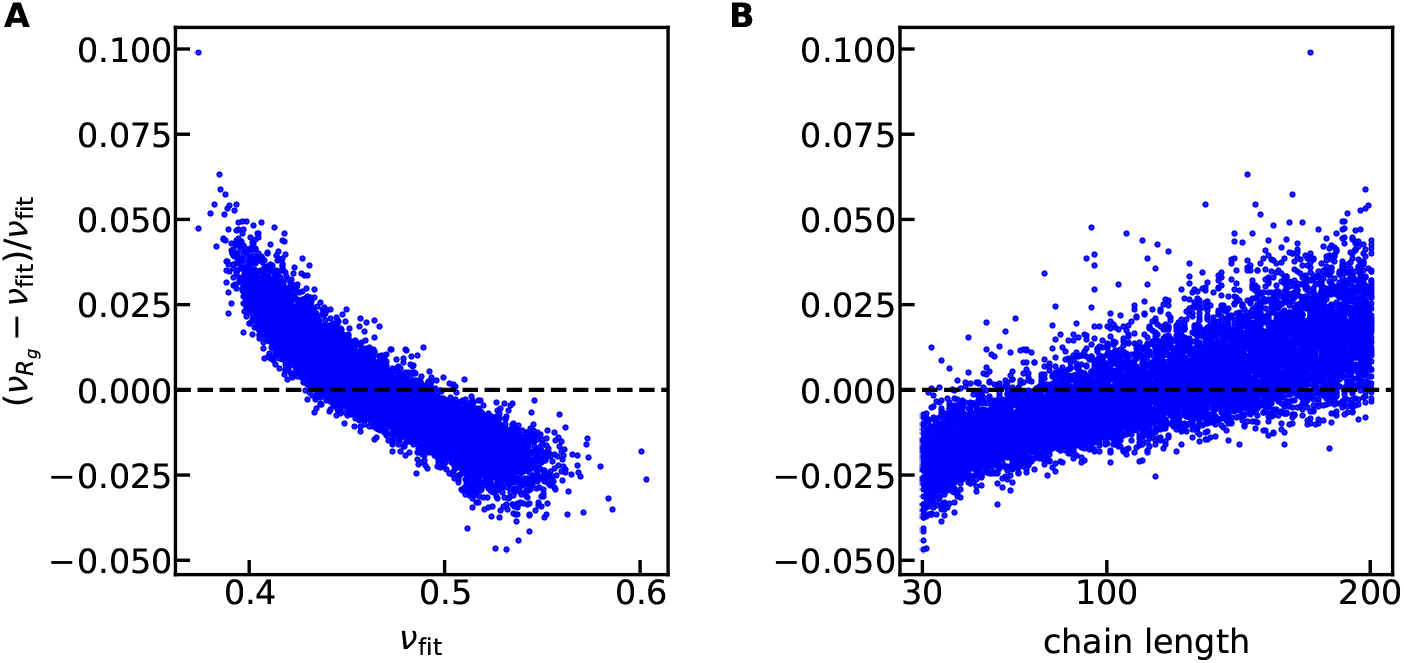
Relative difference between *ν*_Rg_ and *ν*_fit_ as a function of *ν*_fit_ (A) and chain length (B).

**Figure S3.**
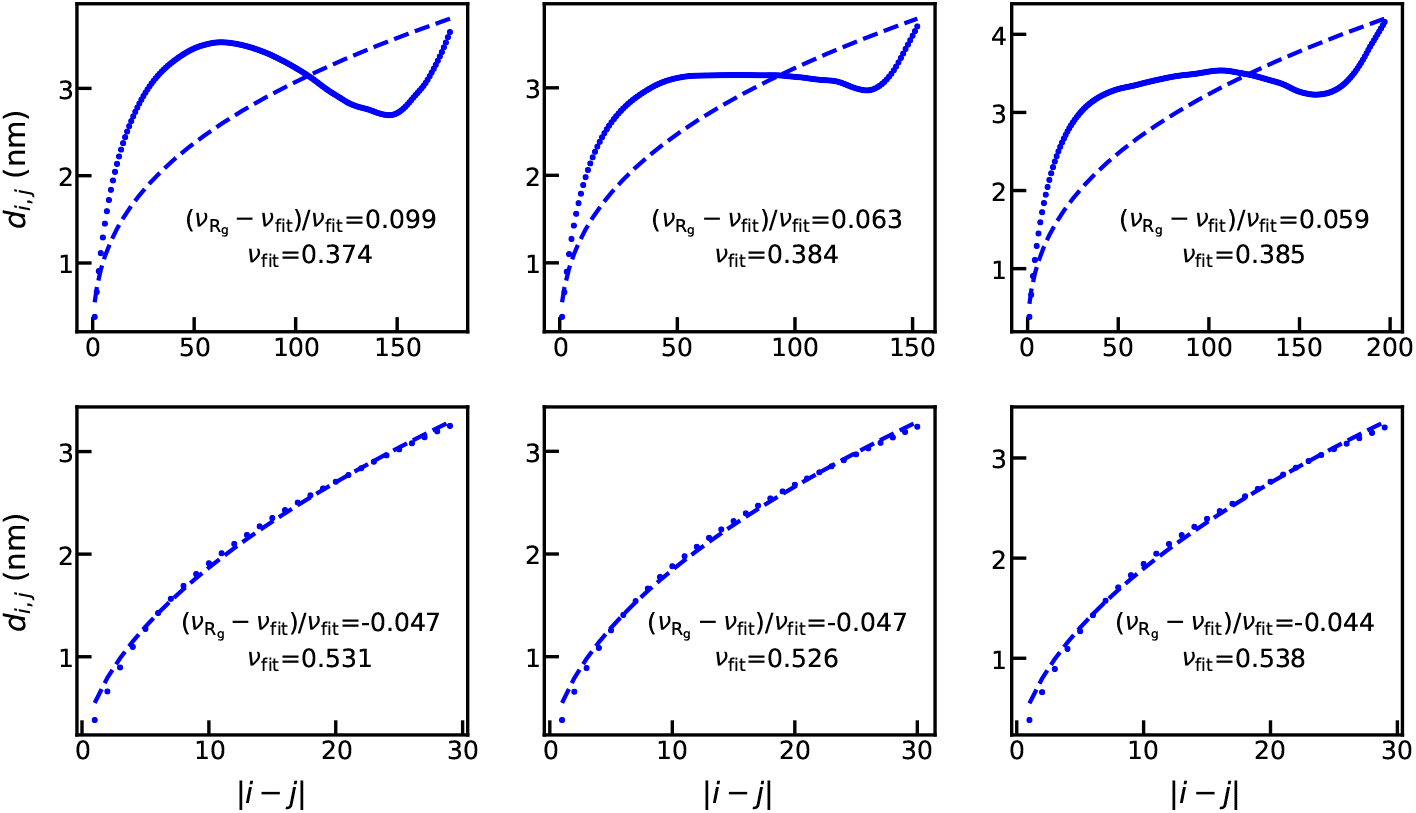
Intrachain distances as a function of the sequence separations for the three cases with most deviations between *ν*_Rg_ and *ν*_fit_: *ν*_Rg_ *> ν*_fit_ (top row) and *ν*_Rg_ *< ν*_fit_ (bottom row). Dotted lines come from simulations and dashed lines show the exponential fitting curves.

**Figure S4.**
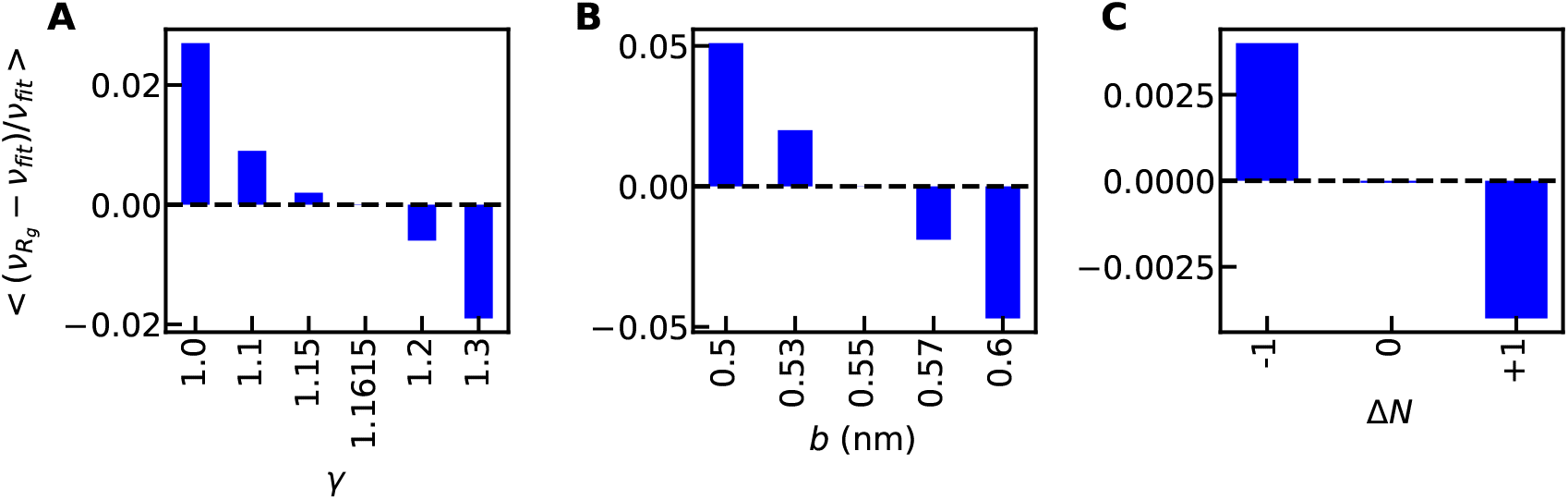
Average relative difference between *ν*_Rg_ and *ν*_fit_ as a function of *γ* (A), *b* (B) and variaton of *N* (C) in Eq. 2 of the main text. Δ*N* = 0 suggests that *N* is defined as the number of peptide bonds and Δ*N* = 1 suggests *N* is defined as the chain length.

**Figure S5.**
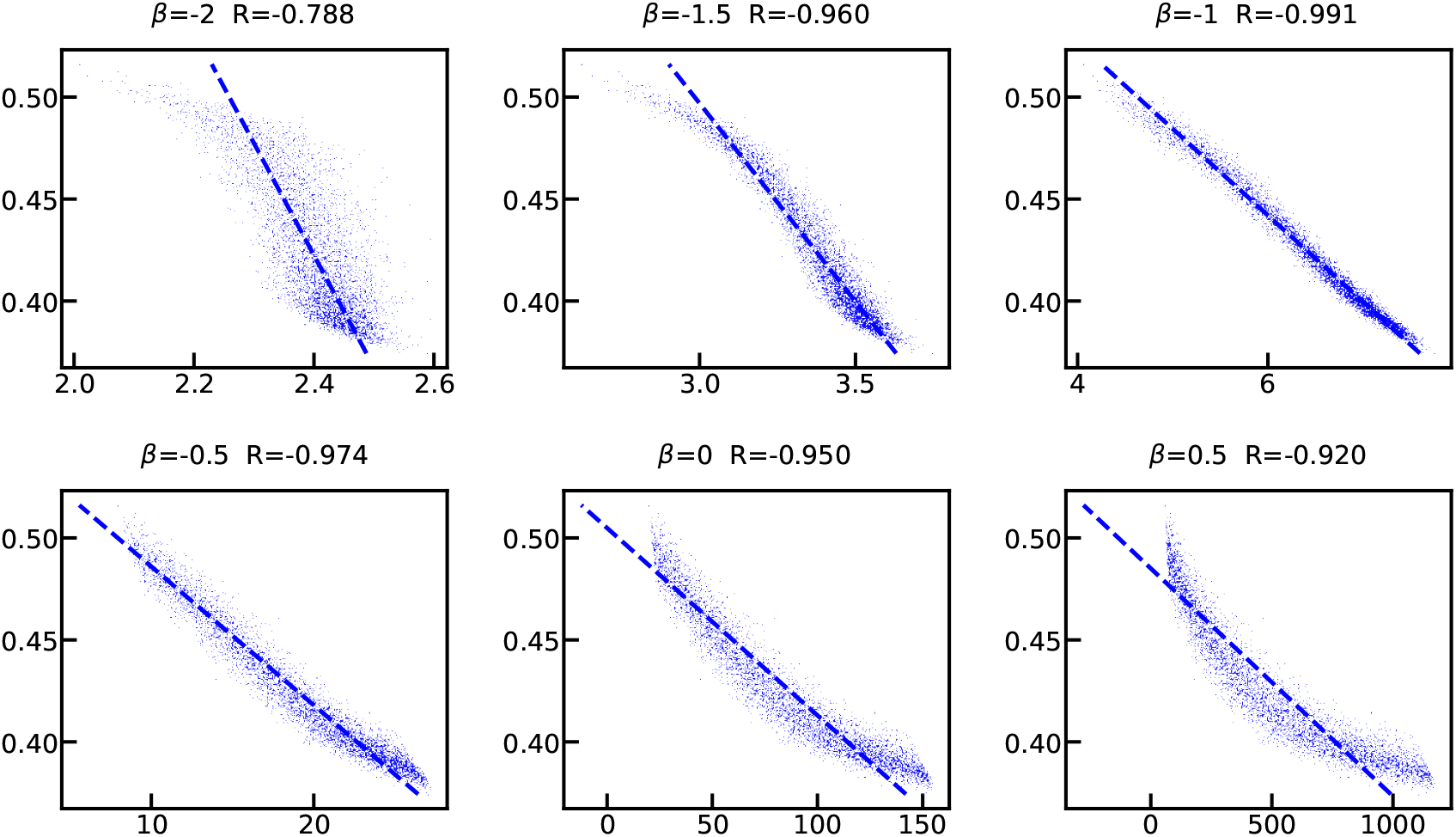
Scanning *β* in defining *SHD* (Eq. 4 of the main text). *R* is the Pearson correlation coefficient between the two variables.

**Figure S6.**
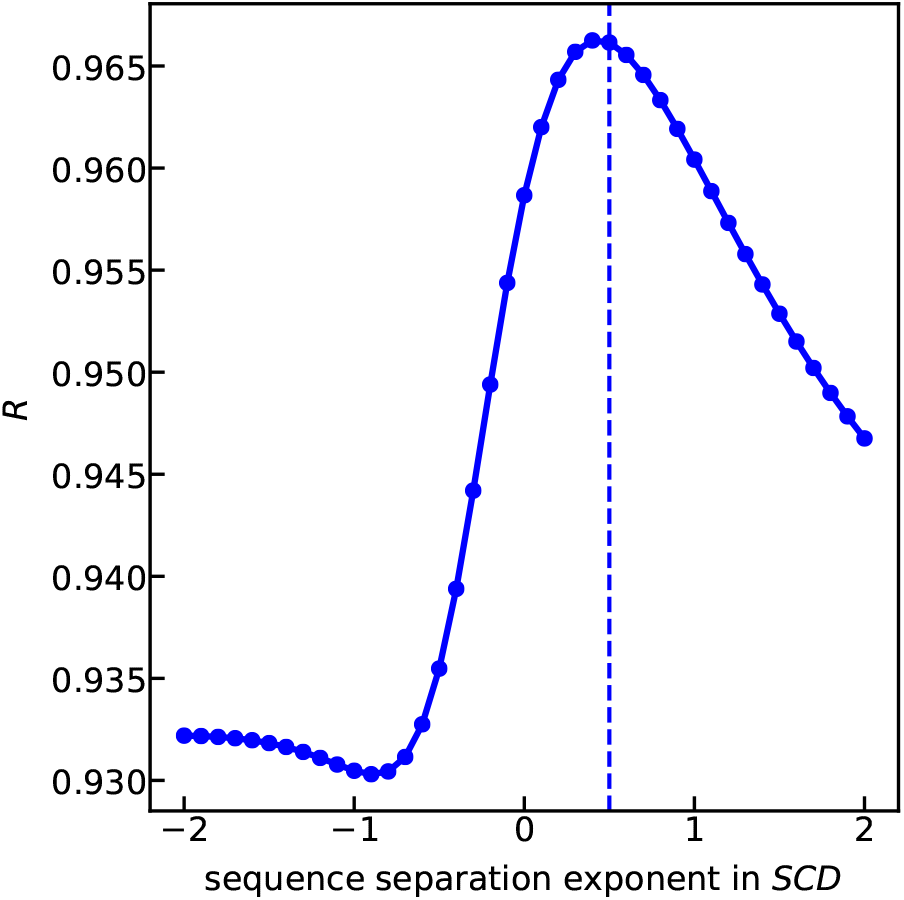
Test of the sequence separation exponent in *SCD*. We use random sequence database A (Fig. S1) for the test. For each scanned exponent in *SCD*, *SHD* is always calculated using an optimal -1 exponent whereas *SCD* is calculated using a different exponent shown in the x-axis. A multilinear regression is then applied to find the best parameters for using *SCD* and *SHD* to fit *ν*. The Pearson correlation coefficient between the modeled and reference *ν* is shown.

**Figure S7.**
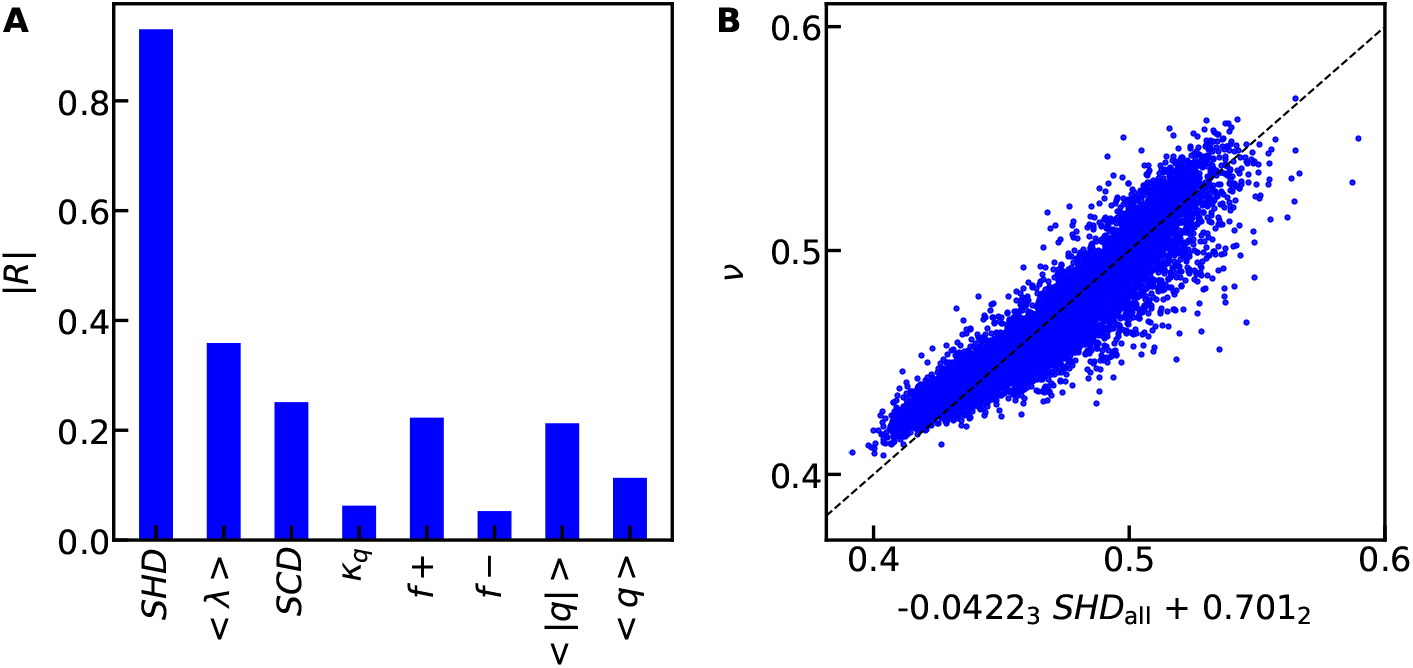
Capturing the scaling exponents using one sequence descriptor. A) Pearson correlation coefficients between the sequence descriptors and simulated scaling exponents. B) The best one sequence-descriptor model with the linear equation shown in labels of x-axis.

**Figure S8.**
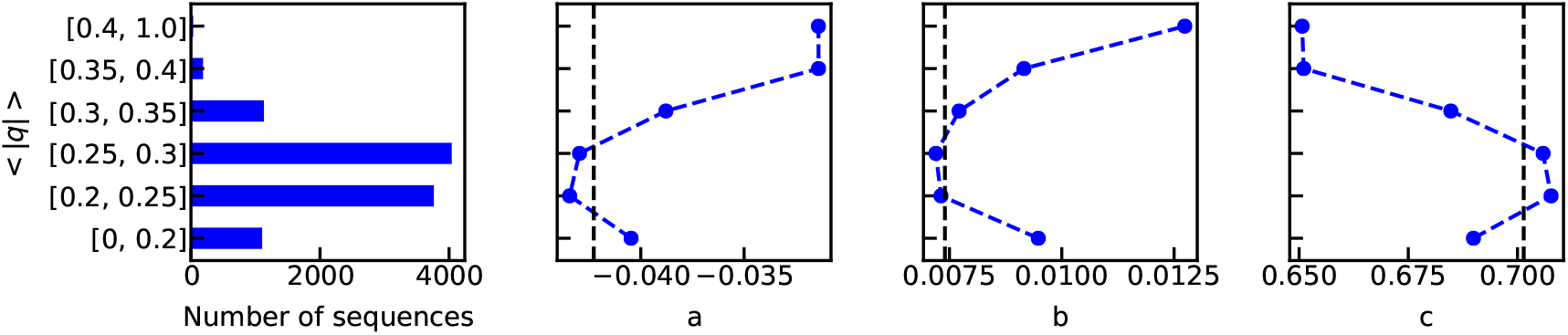
Left: Number of sequences of the database containing charged sequences (Fig. S1) at different range of *< |q| >* values. Right: The three fitting free parameters in the multi-linear equations of *SHD* and *SCD* (Eq. 5 of the main text) when using sequences at different range of *< |q| >* values. The black dashed lines show the fitting parameters using all the sequences.

**Figure S9.**
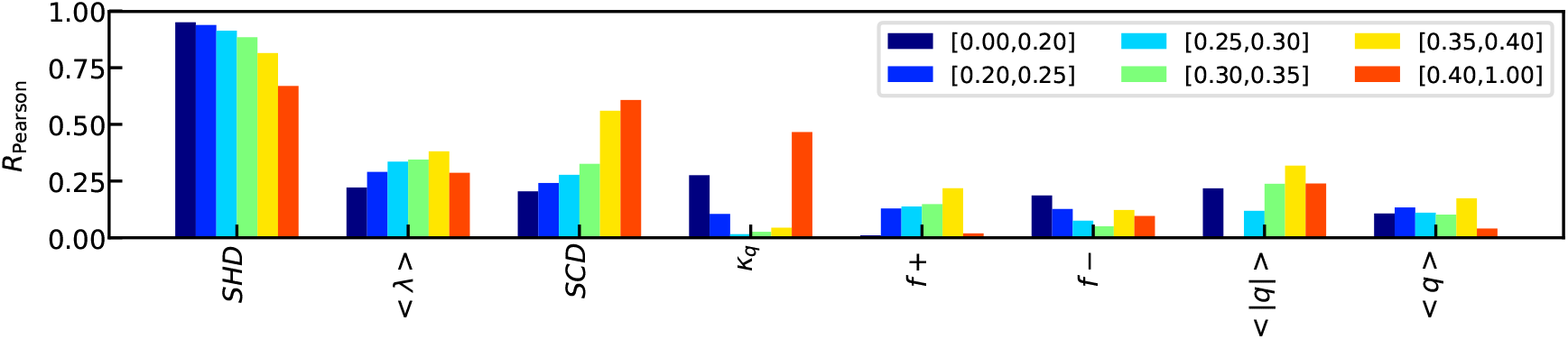
Pearson correlation coefficients between the sequence descriptors and *ν* for sequences with different range of *< |q| >* shown in the legend.

**Figure S10.**
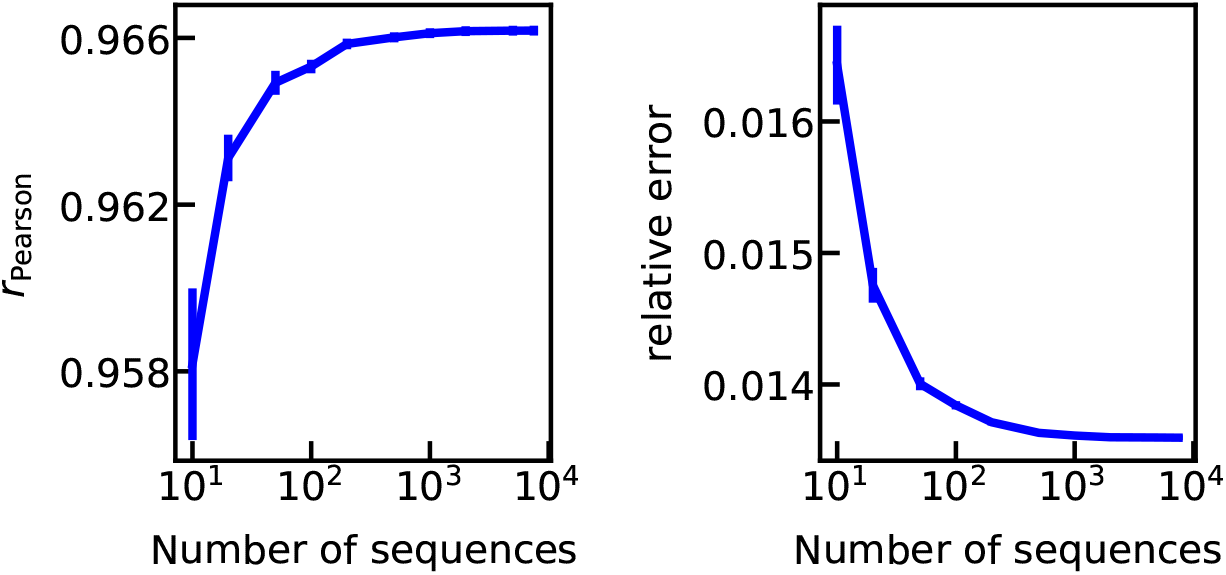
Pearson correlation coefficient (left) and relative errors (right) for assessing the model, when increasing the number of sequences. The error bars are estimated from randomly splitting the database for the training and test sets 100 times.

### 3. Supplementary Tables

**Table S1.**
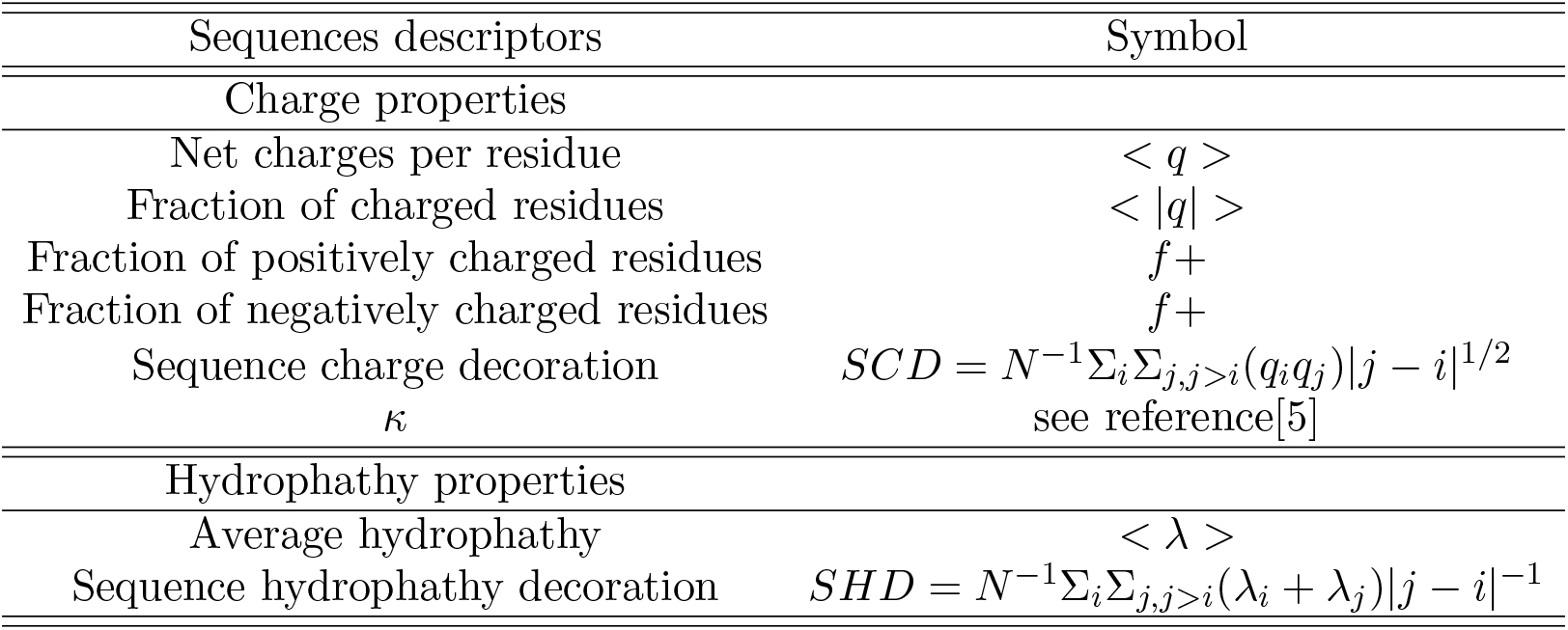
List of sequence descriptors we have tested.

**Table S2.**
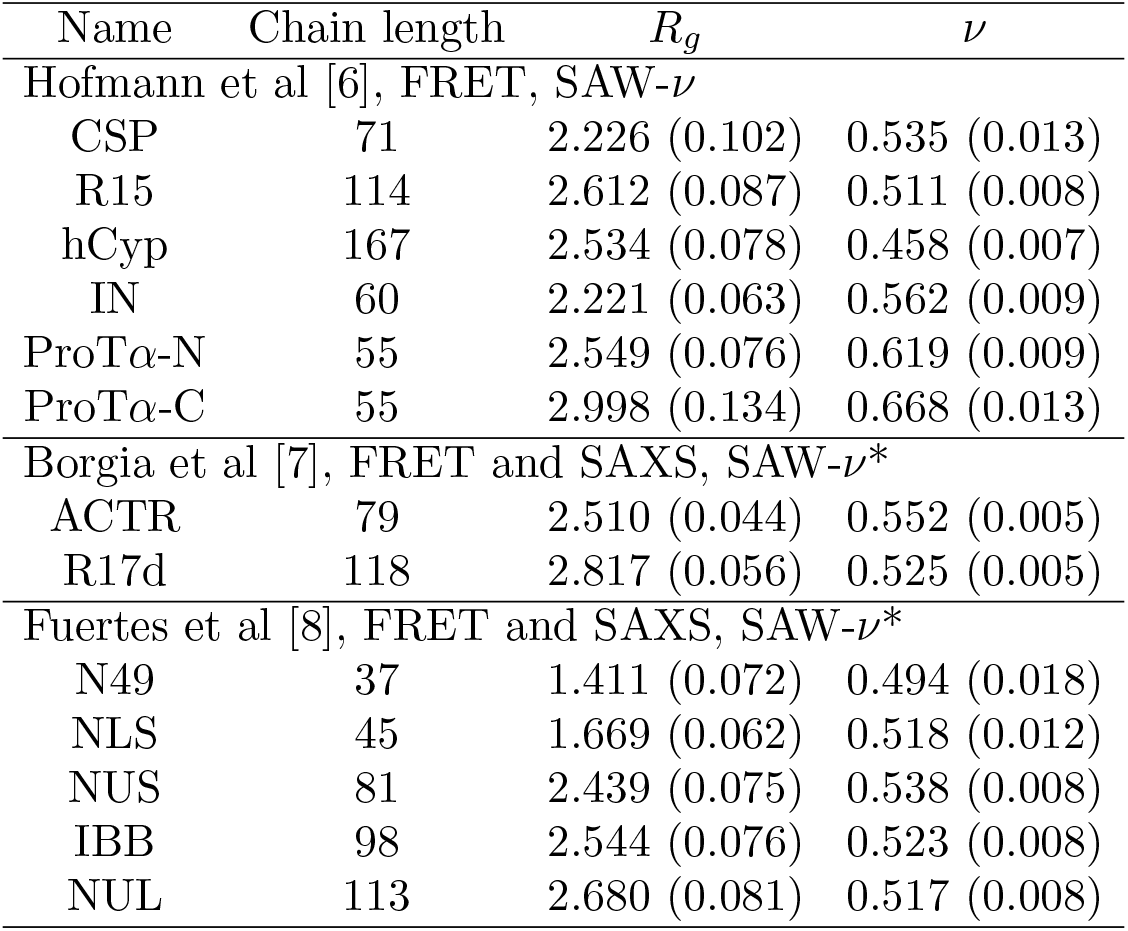
List of sequences with experimentally determined *R_g_*. * Since both FRET and SAXS measurements were provided in these two publications and were shown to be consistent, we only reanalyze the FRET data set here.

